# Genetic mapping of a bioethanol yeast strain reveals new targets for aldehyde- and thermotolerance

**DOI:** 10.1101/2021.11.23.469769

**Authors:** Fellipe da Silveira Bezerra de Mello, Alessandro Luis Venega Coradini, Marcelo Falsarella Carazzolle, Carla Maneira da Silva, Monique Furlan, Gonçalo Amarante Guimarães Pereira, Gleidson Silva Teixeira

## Abstract

Current technology that enables bioethanol production from agricultural biomass imposes harsh conditions for *Saccharomyces cerevisiae*’s metabolism. In this work, the genetic architecture of industrial bioethanol yeast strain SA-1 was evaluated. SA-1 segregant FMY097 was previously described as highly aldehyde resistant and here also as thermotolerant: two important traits for the second-generation industry. A Quantitative Trait Loci (QTL) mapping of 5-hydroxymethylfurfural (HMF) -resistant segregants of hybrid FMY097/BY4742 disclosed a region in chromosome II bearing alleles with uncommon non-synonymous (NS) single nucleotide polymorphisms (SNPs) in FMY097: *MIX23, PKC1, SEA4,* and *SRO77*. Allele swap to susceptible laboratory strain BY4742 revealed that *SEA4*^FMY097^ enhances robustness towards HMF, but the industrial fitness could not be fully recovered. The genetic network arising from the causative genes in the QTL window suggests that intracellular signaling TOR (Target of Rapamycin) and CWI (Cell Wall Integrity) pathways are regulators of this phenotype in FMY097. Because the QTL mapping did not result in one major allelic contribution to the evaluated trait, a background effect in FMY097’s HMF resistance is expected. Quantification of NADPH - cofactor implied in endogenous aldehyde detoxification reactions - supports the former hypothesis, given its high availability in FMY097. Regarding thermotolerance, *SEA4*^FMY097^ grants BY4742 ability to grow in temperatures as high as 38 °C in liquid, while allele *PKC1*^FMY097^ allows growth up to 40 °C in solid medium. Both *SEA4*^FMY097^ and *PKC1*^FMY097^ encode rare NS SNPs, not found in other >1,013 *S. cerevisiae*. Altogether, these findings point towards crucial membrane and stress mediators for yeast robustness.

**KEY POINTS:**

- QTL mapping of the HMF-resistant strain FMY097 reveals a region enriched with SNPs in Chr II
- *SEA4*^FMY097^ has rare non-synonymous mutations and improves cell growth at 10 mM HMF and 38°C
- *PKC1*^FMY097^ has rare non-synonymous mutations and improves cell growth at 40 °C in solid media

## 1. INTRODUCTION

The full exploration of plant potential for fuel or chemical production has become a challenge of great importance in a world with raising environmental awareness. In this matter, lignocellulosic material emerges as the feedstock for second-generation bioethanol (E2G) production, while being vastly available at relatively low cost in different forms (Silveira et al. 2015). The sugarcane E2G industry, one of the most established biorefineries worldwide, is predicated on a separate (SHF) or hybrid (HHF) hydrolysis and fermentation approach (dos Santos et al. 2016). Besides the logistics challenges in implementing industries to produce cellulosic ethanol (*i.e.,* issues related to managing less dense biomass), upstream fermentation processes are complex and still in need of technological improvement. Meanwhile, robust *Saccharomyces cerevisiae* strains are expected to address the bottleneck of ethanol productivity in the currently available operational conditions.

To facilitate enzymatic hydrolysis of cellulose, an acidic thermochemical pretreatment step is required before sugar uptake by *S. cerevisiae*, rendering inhibitory products for the microorganism’s metabolism (Jönsson et al. 2013; Balan 2014). The furan aldehyde 5-hydroxymethylfurfural (HMF), resulting from the dehydration of hexoses - such as D-glucose, D-mannose, D-galactose, and D-fructose (Jayakody et al. 2013) -, is considered one of the most representative inhibitors of *S. cerevisiae* in cellulosic hydrolysates (Nieves et al. 2015). HMF can also further break down to produce levulinic acid, which adds stress to the medium (Liu and Blaschek 2010). *In situ* detoxification of HMF occurs under anaerobic or aerobic conditions in which the aldehyde is reduced in a NAD(P)H-dependent reaction to its less toxic alcohol correspondent – 2,5-bis-hydroxymethylfuran – that is further secreted outside of cells, remaining in the fermentation broth (Liu et al. 2004; Nilsson et al. 2005; Lewis Liu et al. 2008; Liu and Blaschek 2010). Physiologically, HMF is known to increase adaptation time during cultivation, in a dose-dependent manner (Taherzadeh et al. 2000). Furthermore, it has been reported that this aldehyde elevates reactive oxygen species that, generated in mitochondria, induce DNA mutations, protein misfolding and fragmentation, membrane damage, and apoptosis (Almeida et al. 2009). The presence of HMF is also related to redox imbalance that interferes in glycolysis, cell growth, and, finally, biosynthesis (Lewis Liu et al. 2008).

On the other hand, the optimal temperature for cellulase activity ranges from 50 to 55 °C (Pardo and Forchiassin 1999), meaning that a cooling step is demanded prior to microbial fermentation of the sugarcane bagasse hydrolysate. Not only intrinsic process requirements can elevate temperature in industrial mills, climatic circumstances in tropical areas, for instance, can also induce heat stress (Dias et al. 2015) - a remarkable drawback for yeast performance. Therefore, thermotolerance is a desirable trait in industrial strains, given that it could reduce refrigeration costs while increasing ethanol productivity towards a simultaneous saccharification and fermentation (SSF) process (Wang et al. 2019). The effects of heat shock have been deeply revised elsewhere (Yamamoto et al. 2008; Morano et al. 2012; Jarolim et al. 2013), mostly assigning transcription factor Hsf1 and general stress transcription factors Msn2 and Msn4 for regulating thermal response in *S. cerevisiae*. Moreover, thermotolerance is intimately linked to genes involved in the cell wall integrity (CWI) pathway and the actin cytoskeleton (Auesukaree et al. 2009). Also, quantitative trait loci (QTL) mapping has uncovered that alleles *VPS34*, *VID24* and *DAP1* - associated with increased trehalose accumulation or reduced membrane fluidity - have a positive effect on thermal resistance (Wang et al. 2019).

Because microbial robustness towards aldehyde inhibitors and high temperature are crucial in the second-generation industry, in this work we unraveled the genetic basis of these traits in the indigenous bioethanol strain SA-1 (Basso et al. 2008). FMY097 - SA-1-derived haploid, previously described as highly aldehyde tolerant (de Mello et al. 2019) - was used to perform an unprecedented QTL mapping of a population of segregants with elevated HMF resistance. Following, allele replacement in the susceptible BY4742 laboratory strain revealed that *SEA4*^FMY097^ improves growth in presence of HMF. The high availability of NADPH (cofactor implicated in endogenous aldehyde detoxification reactions) in FMY097 suggests that an intricate genetic architecture rules this phenotype in the industrial strain. Further, testing of mutated BY4742 bearing FMY097 alleles in high temperature revealed that *SEA4*^FMY097^ also enhances fitness at 38 °C, while *PKC1*^FMY097^ allowed growth at 40 °C in solid media. Both alleles have rare non-synonymous (NS) mutations that are not present in other >1,013 *S. cerevisiae* strains. These results shed light on the contribution of novel gene targets to important commercial traits in yeast and suggest that signaling transduction pathways TOR (Target of Rapamycin) and CWI are paramount for industrial robustness.

## 2. MATERIALS AND METHODS

### 2.1. STRAINS, PLASMIDS, AND MEDIA

All *S. cerevisiae* strains and plasmids used in this study are described in Table 1. Yeast strains were cultivated in YPD medium (10 g/L yeast extract, 20 g/L peptone, 20 g/L glucose) for propagation or genomic DNA (gDNA) extraction purposes. Geneticin (G418 200 µg/L) was added for the selection of transformed strains with plasmids carrying the *kanMX* drug-resistant marker. Hygromycin (300 µg/mL) was added for the selection of yeast mutants expressing the *hphMX6* cassette. YPD supplemented with 1M sorbitol was used to recover *PKC1* null mutants. YPD cultivations occurred at 30 °C and 250 rpm, when agitation was necessary. Sporulation of diploid cells was performed on 1% KAc medium (1%(v/v) potassium acetate, 460 mg/L of complete drop-out mix) at 25 °C and 250 rpm. For stress screening procedures, yeast strains were cultivated in Synthetic Complete (SC) medium (6.7 g/L yeast nitrogen base, 20 g/L glucose, 460 mg/L of complete drop-out mix). HMF resistance was assessed in SC medium supplemented with either 10 or 20 mM HMF and cultivated at 30 °C, while thermotolerance in SC medium at 38 or 40°C (**see** section 2.11 **for details**). Vector bacteria were grown in Luria-Bertani (LB) broth (10 g/L tryptone, 10 g/L NaCl, 5 g/L yeast extract) supplemented with 100 μg/mL ampicillin at 37 °C and 250 rpm, when agitation was necessary. All media previously described were added 15 g/L of agar for solidification. For long term storage, microorganisms stock solution was kept at −80 °C in media containing 25% glycerol.

**Table 1:**
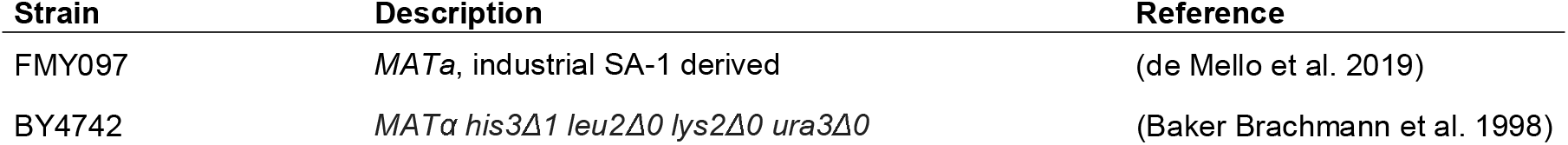
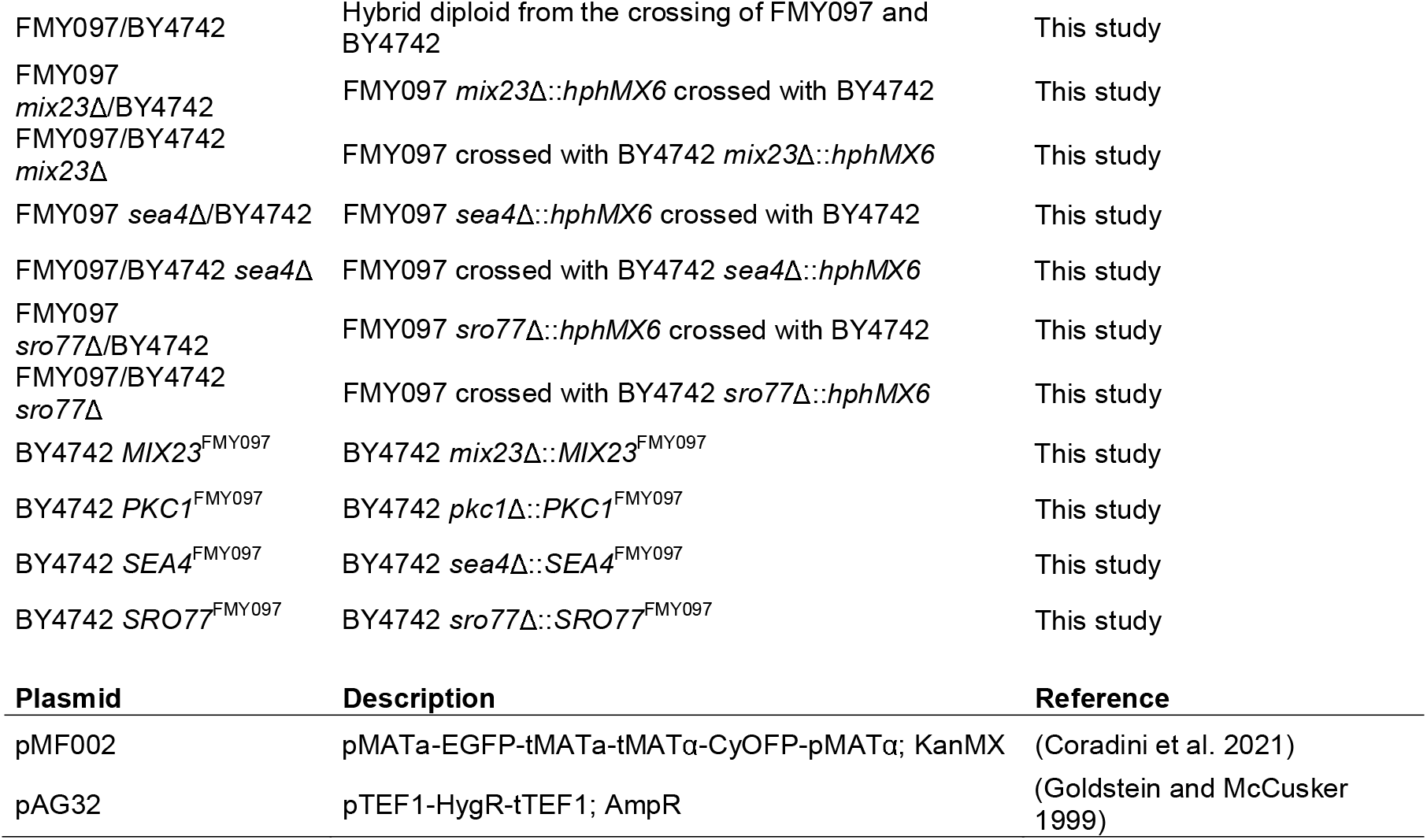
Main *Saccharomyces cerevisiae* strains and plasmids used in this study.

### 2.2. GENERAL MOLECULAR BIOLOGY

For yeast transformation, the LiAc/SS carrier DNA/PEG method (Gietz and Schiestl 2007) was used. Plasmid insertion in DH10β *E. coli* cells was carried out by the traditional electroporation protocol (Dower et al. 1988). Plasmid purification from vector bacteria was performed with the alkaline lysis procedure described by Birnboim and Doly (1979) (Birnboim and Doly 1979). gDNA was extracted with (LiOAc)-SDS/EtOH fast protocol (Lõoke et al. 2017) for polymerase chain reaction (PCR) purposes or according to the procedures described by Ausubel et al. (1988) (Ausubel et al. 1998) for sequencing. PCR reactions were performed with Phusion High Fidelity DNA Polymerase following the manufacturer’s instructions (NEB – New England Biolabs). All primers used in this study are available in **Supplementary Table S1**.

### 2.3. PARENTAL DIPLOID CONSTRUCTION AND SPORULATION FOR QTL MAPPING

For the QTL mapping of HMF resistance, the highly heterozygous hybrid was obtained by crossing the strains FMY097 (HMF resistant) and BY4742 (HMF susceptible). The mating procedure concerned over-scratching both strains in a YPD plate. After overnight incubation at 30°C, single colonies were isolated, and their mating type was PCR-checked. The confirmed F_1_ diploid FMY097/BY4742 was transformed with plasmid pMF002 for further haploid collection, as described by (Coradini et al. 2021). Sporulation was performed on conical glass tubes containing 2 mL of 1%KAc medium for 5-7 days.

### 2.4. F_2_ SPORES DISRUPTION

For high-throughput haploid isolation using a flow cytometry-based approach, FMY097/BY4742 spores were first treated for *asci* digestion. Initially, 250 µL of the spore culture was centrifuged in an Eppendorf tube, and the cell pellet was resuspended in 100 µL of micromanipulation buffer, containing 1 µL of β-mercaptanol. Following, 16 µL of cells added with 20 µL Lyticase (0.5 mg/mL) was incubated at 30°C and 900 rpm for three hours. *Asci* digestion was microscope-inspected and interrupted with 100 µL of distilled water. Cell suspension was then washed 3 times with distilled water. Finally, 1 mL Triton (0.01%) was added to the test tube and sonicated (Fine Point model) at level 2 (20%) for 1 minute - this process was repeated, and Eppendorf tubes were kept in ice. 250 µL of cell suspension was plated in YPD + G418 and cultivated for 2 days at 30°C.

### 2.5. FLOW CYTOMETRY F_2_ SEGREGANTS COLLECTION

To isolate FMY097/BY4742 haploid segregants from other diploid or sporulated cells, colonies grown in the plate after *asci* disruption were collected and sorted in flow cytometry. Initially, hypothetical haploid plates were washed with 5 mL of 1X PBS and collected in a 50 mL falcon. The suspension was sonicated twice for 1 minute at level 2 (20%) and then diluted in 1X PBS to optical density (OD_600_) = 0.4. High sensitivity flow cytometer BD FACSARIAS III (BD-Bioscience) was used to sort haploid yeast, as follows: 500,000 green (MATa, 515/545 nm) or orange (MATα, 655/695 nm) fluorescent cells each, using a FITC filter (488 nm). The collected cells were cultivated in 10 mL YPD containing 100 µg/mL ampicillin in glass tubes for 6 hours at 30 °C. Cultures were centrifuged and resuspended in 950 µL water. 200 µL of cell suspension were then plated on YPD and grown overnight at 30°C. Finally, individual colonies were transferred to 150 µL YPD medium in 96-well flat bottom plates. The ploidy of the segregants was confirmed by mating type testing using the halo approach described by Julius et. al (Julius et al. 1983).

### 2.6. PHENOTYPING OF HAPLOIDS LIBRARY IN HMF AND POOL SELECTION

Phenotyping of segregants was performed as described by Matsui et. al (2016) (Matsui and Ehrenreich 2016). FMY097/BY4742 haploids stock solution in microtiter plates were inoculated on 96-well plates containing 200 µL of YPD using a 96-pin replicator and let grow overnight at 30 °C. After reaching stationary phase, the plates were agitated for 30 seconds using a plate spectrophotometer (Spectramax Plus 384) for homogenization and segregants were further pinned on SC plates containing 20mM HMF. The strain’s positions were randomized in triplicates in order to minimize technical errors. Growth of each strain colony spot was assessed after 48h incubation by plate image capture using Gel Doc XR+ Gel Documentation System (BIO-RAD). The dimensions of all the images were set at 13.4 x 10 cm (W x L) and captured under white Epi illumination with an exposure time of 0.5 seconds. The pixel intensity of each colony was measured by ImageJ software. The total pixel intensity within a circle (spot radius = 50 pixels) surrounding each colony in the image was measured using the *Plate Analysis JRU v1* plugin for ImageJ. Average pixel intensity - determined by dividing the total pixel intensity by the area of the circle examined (7845 pixels^2^) - was used as the parameter for HMF resistance. Relative growth was then calculated as the ratio of pixel intensity observed in the treatment and control conditions. Haploids that presented relative colony size with a Z-score > 1.5 were selected as individuals with extreme HMF resistance. 60 superior segregants (displaying high tolerance to HMF) were organized in the “BEST” pool, while inferior segregants (susceptible to 20 mM HMF) were assembled in the “WORST” pool.

### 2.7. PREPARATION OF THE DNA SAMPLE

The 60 superior and inferior segregants pools for the HMF resistant QTL mapping and parental BY4742 were individually inoculated in 2 mL YPD medium and grown overnight at 30°C. gDNA was pooled-extracted using the method described by Pais et al. (2014) (Pais et al. 2014). Initially, OD_600_ of each strain in the pool was individually measured and cells mixed in equivalent concentrations in order to form a heterogeneous culture containing approximately the same quantity of cells representative of each strain in the pool. Following gDNA extraction, DNA concentration and quality were assessed with the Nanodrop 3000 UV-Vis spectrophotometer (Wilmington). FMY097’s genome sequence was already available (Nagamatsu et al. 2019).

### 2.8. POOLED-SEGREGANT WHOLE-GENOME SEQUENCE ANALYSIS

Aliquots of gDNA from the segregant pools and parental BY4742 containing at 5 μg were provided to Life Sciences Core Facility (LaCTAD) from the University of Campinas (UNICAMP) for whole-genome sequencing using the Illumina HiSeq 4000 platform. The generated 2×100 paired-end reads were initially aligned to the genome sequence of CEN.PK113-7D (Nijkamp et al. 2012) using program Bowtie2 version 2.3.5.1 (Langmead and Salzberg 2012). The alignment was then converted to BAM files using samtools software version 1.3.1 (Li et al. 2009). When multiple read pairs presented identical external coordinates, the pair with the highest mapping quality was retained using picard version 2.23.9 command “MarkDuplicates” (https://broadinstitute.github.io/picard/). SNP calling was performed using GATK (v4.0.12.0) base quality score recalibration, indel realignment and SNP and INDEL discovery and genotyping across the two samples (“BEST” and “WORST” pool) using standard hard filtering parameters (McKenna et al. 2010). For each pool the ploidy level was configured to 60 (number of haploid individuals in each pool).

### 2.9. QTL MAPPING

QTL Bulk Segregant Analysis (BSA) of HMF resistance was performed based on the distribution of SNP-index and Δ(SNP-index) over the chromosomes, according to Takagi et al. (2013) (Takagi et al. 2013). SNP-index was calculated by dividing the number of the alternative variant by the total number of aligned reads. The analysis is based on calculating the allele frequency differences, or Δ(SNP-index), from the allele depths at each SNP. In this work, SNP-index value was 0 when all short reads contain genomic fragments from BY4742, and SNP-index value was 1 when all short reads contain SNPs from the other parental FMY097. The difference between the SNP-index of *“BEST* pool” and that of “*WORST* pool” was calculated as the Δ(SNP-index). A very high Δ(SNP-index) was indicative of a one-sided SNP preferentially inherited from the superior parental, indicating a genetic linkage to high tolerance to HMF. Additionally, according to the SNP density, a sliding window analysis with 50 kb window size was used for the calculation of the average Δ(SNP-index) of SNPs located in a certain genomic interval, and the extreme quantiles are used as simulated confidence intervals (CI) of 99% or 95%. Δ(SNP-index) graph was plotted by aligning an average Δ(SNP-index) against the position of each sliding window in the genome. Regions that surpass the CI of 0.99 were considered putative QTLs. Simulations and calculations were performed using the R package *QTLseqr* (Mansfeld and Grumet 2018).

### 2.10. GENETIC ENGINEERING

Genetic modification of strains was performed using a CRISPR-Cas9 system. Reciprocal hemizygotes were obtained from the crossing of parental BY4742 or FMY097 with a given allele knocked out and the non-edited remaining strain. For gene knock-out, the transformation was carried out with a Cas9+sgRNA plasmid containing a specific single-guided RNA (sgRNA) sequence homologous to the target region and a double-stranded DNA (dsDNA) donor PCR-amplified from plasmid pAG32, bearing the hygromycin resistance cassette *hphMX6* and flanking regions homologous to the external region of the target gene open reading frame (ORF). Null mutant strains were collected in hygromycin plates. Allele interchange was performed on the knocked-out versions of strain BY4742. For that, the CRISPR-Cas9 plasmid was constructed with a sgRNA homologous to *hphMX6* and the donor dsDNA consisted of the ORF PCR-amplified from FMY097’s genome. Mutants were confirmed via PCR or Sanger sequencing.

### 2.11. CHARACTERIZATION OF STRESS TOLERANCE

Screening of HMF resistance and thermotolerance traits of the strains was performed in both solid and liquid media. For solid media, yeasts were inoculated in agar plates in serial dilutions (starting from OD_600_ = 1) and photographed after 72 h incubation. Phenotyping in liquid media was assessed in static micro cultivation in 96-well plates, with an initial OD_600_ of 0.1 in a total volume of 150 µL (135 µL medium + 15 µL inoculum with OD_600_ = 1). All cultivations were performed using 4 replicates. Optical density data were collected every 1 hour for 72 h total using a plate reader (SpectraMAX 384, Molecular Devices). Growth kinetic parameters were calculated using the MATLAB code OCHT reported by de Mello *et al.* (2019) (de Mello et al. 2019). Statistical analysis was performed using Student’s t-test.

## 3. RESULTS

### 3.1. SEGREGANT LIBRARY COLLECTION OF A BIOETHANOL HYBRID STRAIN

Given that the acid pretreatment of lignocellulosic feedstock usually yields inhibitory aldehyde compounds and that the biomass hydrolysis requires high temperature for enzymatic activity, *S. cerevisiae* strains that present robustness towards both fermentation challenges are good candidates for SSF processes. Indigenous bioethanol yeasts have been praised for their fermentation performance under harsh conditions, in special Brazilian E1G strains PE-2, SA-1, and others (Basso et al. 2008). SA-1 haploid segregant FMY097 has been previously described as highly aldehyde resistant and suggested as a good chassis for the second-generation industry (de Mello et al. 2019).

Here, FMY097 and laboratory BY4742 were phenotyped in presence of 20 mM HMF and at 40 °C in both liquid and solid media (Fig 1a), in order to assess the strain’s response to both cultivation stresses. Growth in liquid media allowed the analysis of final culture cell density, Final OD (Fig 1c and 1f), and the calculation of the maximum specific growth rate, µ_MAX_ (Fig 1d and 1g). All data concerning the kinetic growth parameters are described in Table 2 as well.

**Fig 1:**
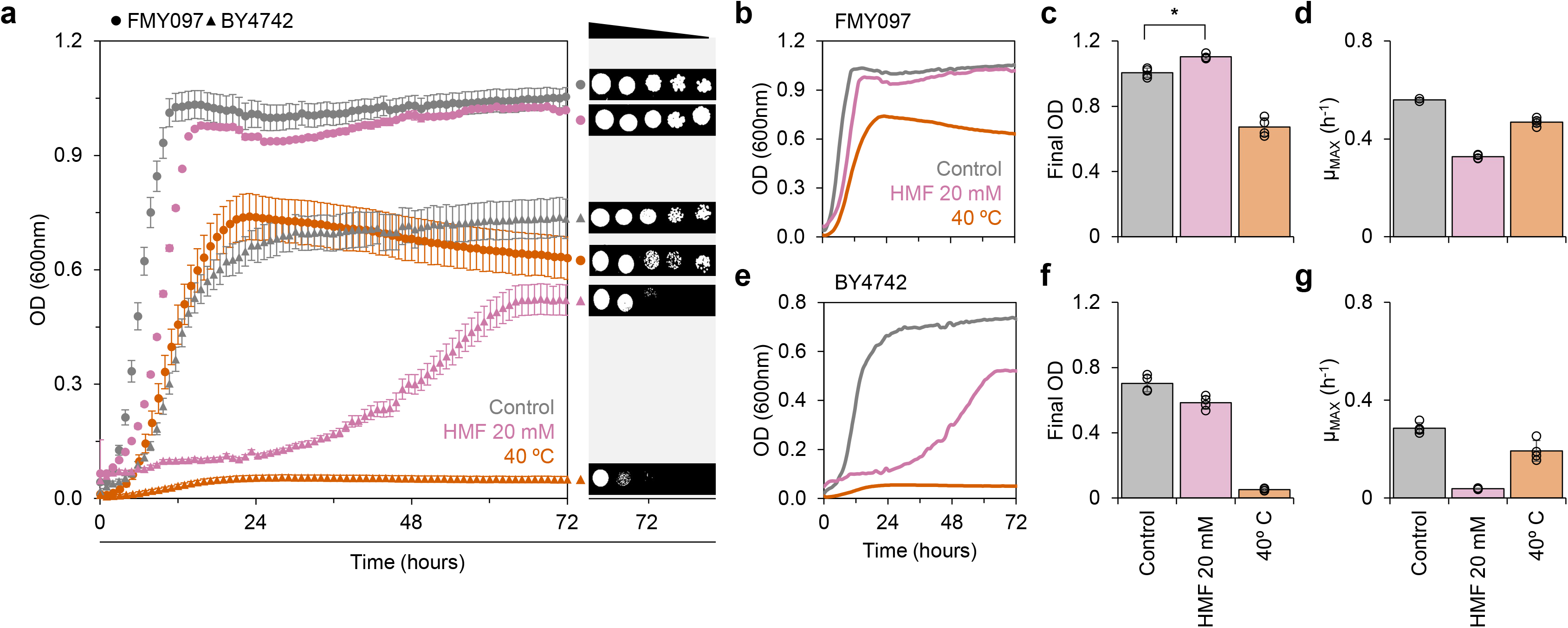
Phenotyping of strains FMY097 (MATa, industrial SA-1 derived) and BY4742 (MAT) in HMF and high temperature. **a:** *On the left*: growth curve of 4 replicates of strains FMY097 (triangle) and BY4742 (circle) in SC 30 °C (control, grey), SC + 20 mM HMF 30°C (pink) and SC 40 °C (orange); *on the right*: spot test of serial dilutions of strains in the same conditions after 72h incubation - each image is labeled with the corresponding format (strain) and color (medium) on the left; **b and e:** representative growth curves of strains FMY097 and BY4742, respectively, in the same conditions; **c and f:** bar charts of the final OD of strains FMY097 and BY4742, respectively; **d and g:** bar charts of the maximum specific growth rate of strains FMY097 and BY4742, respectively. Black circles in the bar charts represent the replicate data. (*) stands for p-value < 0.05

**Table 2:** Kinetic growth parameters of the main strains in different cultivation conditions.

|  | Control | HMF 10 mM | HMF 20 mM | 38°C | 40°C |
| --- | --- | --- | --- | --- | --- |
| Strain | Final OD (600nm) |  |  |  |  |
| FMY097 | 1.005 ± 0.028 | 1.124 ± 0.028 | 1.104 ± 0.015 | 0.669 ± 0.047 | 0.673 ± 0.056 |
| BY4742 | 0.702 ± 0.050 | 0.535 ± 0.084 | 0.585 ± 0.040 | 0.334 ± 0.042 | 0.052 ± 0.008 |
| BY4742 MIX23<sup>FMY097</sup> | 0.711 ± 0.015 | 0.600 ± 0.057 | 0.365 ± 0.061 | 0.375 ± 0.032 | 0.051 ± 0.005 |
| BY4742 *PKC1*<sup>FMY097</sup> | 0.727 ± 0.022 | 0.430 ± 0.023 | 0.309 ± 0.058 | 0.292 ± 0.019 | 0.041 ± 0.022 |
| BY4742 *SEA4*<sup>FMY097</sup> | 0.716 ± 0.093 | 0.672 ± 0.020* | 0.404 ± 0.136 | 0.442 ± 0.044* | 0.034 ± 0.005 |
| BY4742 *SRO77*<sup>FMY097</sup> | 0.655 ± 0.075 | 0.462 ± 0.044 | 0.323 ± 0.025 | 0.386 ± 0.046 | 0.024 ± 0.000 |
| *Strain* | $\mu_{\text{MAX}}$ (h<sup>-1</sup>) | | | | |
| FMY097 | 0.699 ± 0.243 | 0.408 ± 0.018 | 0.327 ± 0.008 | 0.489 ± 0.014 | 0.468 ± 0.017 |
| BY4742 | 0.286 ± 0.022 | 0.124 ± 0.012 | 0.038 ± 0.002 | 0.180 ± 0.004 | 0.193 ± 0.043 |
| BY4742 *MIX23*<sup>FMY097</sup> | 0.297 ± 0.023 | 0.102 ± 0.013 | 0.036 ± 0.015 | 0.217 ± 0.005*** | 0.208 ± 0.034 |
| BY4742 *PKC1*<sup>FMY097</sup> | 0.244 ± 0.017 | 0.137 ± 0.016 | 0.047 ± 0.009 | 0.189 ± 0.011 | 0.209 ± 0.071 |
| BY4742 *SEA4*<sup>FMY097</sup> | 0.268 ± 0.022 | 0.094 ± 0.008 | 0.021 ± 0.005 | 0.202 ± 0.005*** | 0.208 ± 0.051 |
| BY4742 *SRO77*<sup>FMY097</sup> | 0.263 ± 0.021 | 0.093 ± 0.006 | 0.029 ± 0.008 | 0.215 ± 0.018*** | - |

Asterisks represent statistical difference observed between mutated BY4742 and parental FMY097. (\*) stands for p-value < 0.05 and (\*\*\*) for p-value < 0.001.

This initial assay confirmed FMY097’s HMF resistance and validated its thermotolerance. In fact, the strain was able to outgrow the control condition when 20 mM HMF was present in the medium, reaching OD_600_ of 1.104 ± 0.015 in contrast to 1.005 ± 0.028. Concerning growth at 40 °C, FMY097 showed a robust response, with little compromising of its µ_MAX_ - *i.e.,* the strain presented fast growth but lost 33% of cell production. These results reassert that strain FMY097 should be a viable background for industrial applications. On the other hand, laboratory BY4742 growth was significantly impaired in both conditions. Although 20 mM HMF did not fully repress BY4742 growth, yielding a final OD_600_ of 0.585 ± 0.040, a long adaptation time can be observed. Meanwhile, 40 °C was a limiting condition for the laboratory strain, representing a lethal environment with no cell growth.

Because the QTL approach to uncover the genetic basis of aldehyde resistance in yeast has not yet been described, and FMY097 has been reported to resist up to 80 mM HMF (de Mello et al. 2019), we moved towards this analysis bearing in mind that this is also a thermotolerant chassis. A visual summary of the methodology used in this work for obtaining a genetically diverse segregant population, regarding their HMF tolerance, is presented in **Supplementary Fig S1**. Initially, we crossed the strains FMY097 and BY4742, resulting in the heterozygous F_1_ hybrid FMY097/BY4742. The resulting diploid was transformed with plasmid pMF002 expressing different gene reporters with specific mating-type promoters, and further sporulated, as described by Coradini *et al* (Coradini et al. 2021). Following the *asci* lysis, haploid collection was performed using flow cytometry. A total of 952 hypothetical haploids were collected, being 472 “MATa” and 480 “MATα”. Confirmation of ploidy by the halo approach confirmed that from the MATa population, 37 were diploids and the remaining 435 were haploids indeed (being 434 MATa and 1 MATα) – 92% efficiency in the correct mating type selection. Out of the 480 “MATα” collected organisms, 421 were diploids and only 59 haploids (being 21 MATa and 38 MATα) – collection efficiency of 8%. Therefore, a library of 494 haploid segregants from the FMY097/BY4742 hybrid was obtained – enough organisms for the QTL analysis.

### 3.2. PHENOTYPING OF THE HAPLOID SEGREGANTS COLLECTION

Phenotyping of the FMY097/BY4742 segregant library was performed as described by Matsui *et. al* (Matsui and Ehrenreich 2016): in short, HMF-resistance was assessed as a function of colony size in solid media containing HMF. A concentration of 20 mM HMF was set as the minimal inhibitory concentration (*i.e.,* the minimal concentration capable of inhibiting the growth of at least one segregant - data not shown). After 48h of cultivation at 30 °C, the plates were photographed and each segregant’s colony size was determined according to their pixel area. A normal distribution of colony size throughout the population was observed, confirming that HMF resistance is a quantitative trait. The Z-score value for each segregant was further calculated and 93 haploids with Z > 1.5 were selected for a new round of phenotyping (**see Supplementary Fig S2**).

For the re-phenotyping, each haploid colony size was assessed in triplicates using the random block approach. Out of the 93 segregants, 60 were selected with a p-value < 0.05 relative to the hybrid parental FMY097/BY4742 performance. This group of haploids was named “*BEST*” pool. Other 60 segregants, which presented the lowest values for colony size in the first round of phenotyping – *i.e.,* non-resistant haploids – were grouped and labeled “*WORST*” pool. Inferior segregants pools are necessary as a control group in the QTL mapping. Colony size distribution of the segregants in the *BEST* and *WORST* pools are very distinct. For the first, colony size (pixels) average is 48.73 ± 3.08 and, for the last, 38.58 ± 2.08 (p-value < 0.001) (**see Supplementary Fig S3**). The clear phenotypic difference regarding the HMF resistance between the two pools guarantees a precise mapping of the genomic regions in which there are polymorphisms comprising a QTL region.

### 3.3. QTL MAPPING OF HMF RESISTANCE IN *S. cerevisiae*

Genomic DNA from both strain BY4742 and the samples “*BEST* pool” and “*WORST* pool” were subjected to whole-genome sequencing analysis using the Illumina HiSeq 4000 platform. FMY097 genome has been previously reported (Nagamatsu et al. 2019). To identify SNPs, the sequence reads from the parental strains were first aligned to the CEN.PK113-7D reference genome sequence. 38820 SNPs were identified between BY4742 and FMY097 and further selected for QTL analysis.

Using *QTLseqr* (Mansfeld and Grumet 2018), the Δ(SNP-index) was calculated and a sliding window analysis with 50 kb size and 200 bp increment was used for the calculation of the average Δ(SNP-index) and confidence interval (CI) (99% or 95%) of SNPs located in the same genomic region. A Δ(SNP-index) graph was plotted by aligning an average Δ(SNP-index) with the position of each sliding window in the genome (Fig 2). For this study, regions above a CI of 0.99 (blue line) were considered candidates for major contribution to the phenotype. One genomic region in chromosome II with Δ(SNP-index) values above CI of 0.99 from 2867 to 40972 bp was identified. At least 3 other *loci* (on chromosomes IX, XII, and XIII) with high values of Δ(SNP-index) are visible but did not make the established threshold and were not considered. Although there are peak regions of negative Δ(SNP-index) close to the CI of 99% (on chromosomes IV, XIII, and XV), these weren’t selected as possible QTL regions given their relation to the genomic polymorphisms of the low-tolerant segregants pool (*i.e.,* mutations that do not confer HMF resistance).

**Fig 2:**
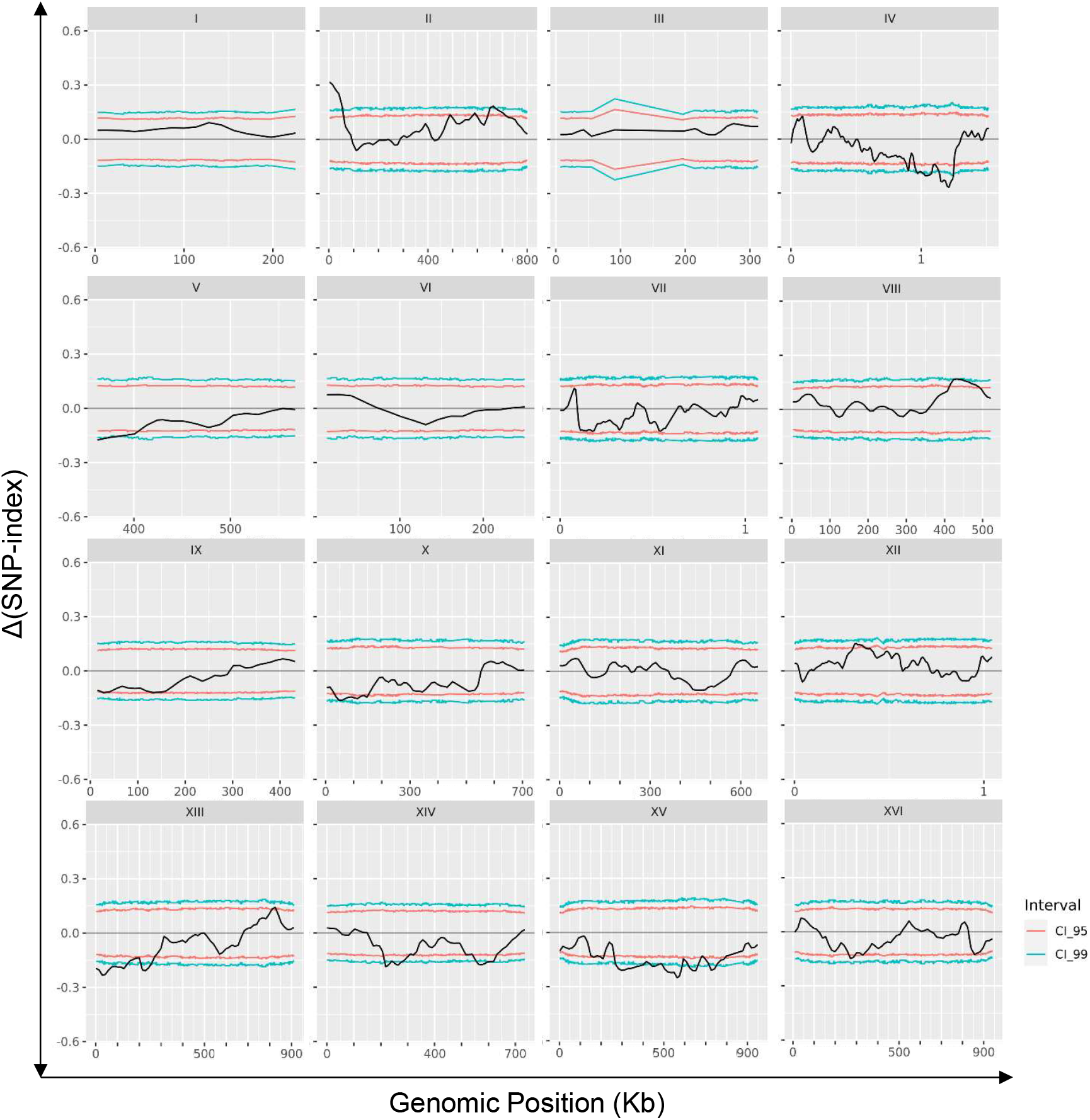
QTL mapping of HMF resistance: mapping of the *loci* involved in the trait by pooled-segregant whole-genome sequence analysis. The X-axis indicates the chromosome’s position; Y-axis indicates the Δ(SNP-index) values. Δ(SNP-index) is the difference between the SNP-index of “*BEST* pool” and that of “*WORST* pool.” Black lines represent the average Δ(SNP-index) values, as determined using a sliding window analysis with 50 kb window size and 200 bp increment. Red lines represent the confidence interval of 95% and blue dotted line, 99% - used threshold for putative regions responsible for HMF resistance

A total of 29 coding sequences are present in the candidate region located on chromosome II. Detailed analysis of the FMY097 genome sequence of this region revealed that 9 alleles present mutations (**see Supplementary Table S2**). Out of these, 6 alleles contain at least one non-synonymous SNP in the open reading frame. The prevalence of these SNPs in other *S. cerevisiae* strains was checked to eliminate mutations that might not be related to the HMF resistance phenotype. The disposition of NS mutations in the alleles observed in the QTL region in chromosome II among other *S. cerevisiae* strains is available in **Supplementary Table S3**. Finally, we narrowed it down to alleles *MIX23*, *PKC1*, *SEA4,* and *SRO77* (Table 3) bearing at least one non-synonymous uncommon mutation in the superior parental FMY097 for further functional analysis.

**Table 3.**
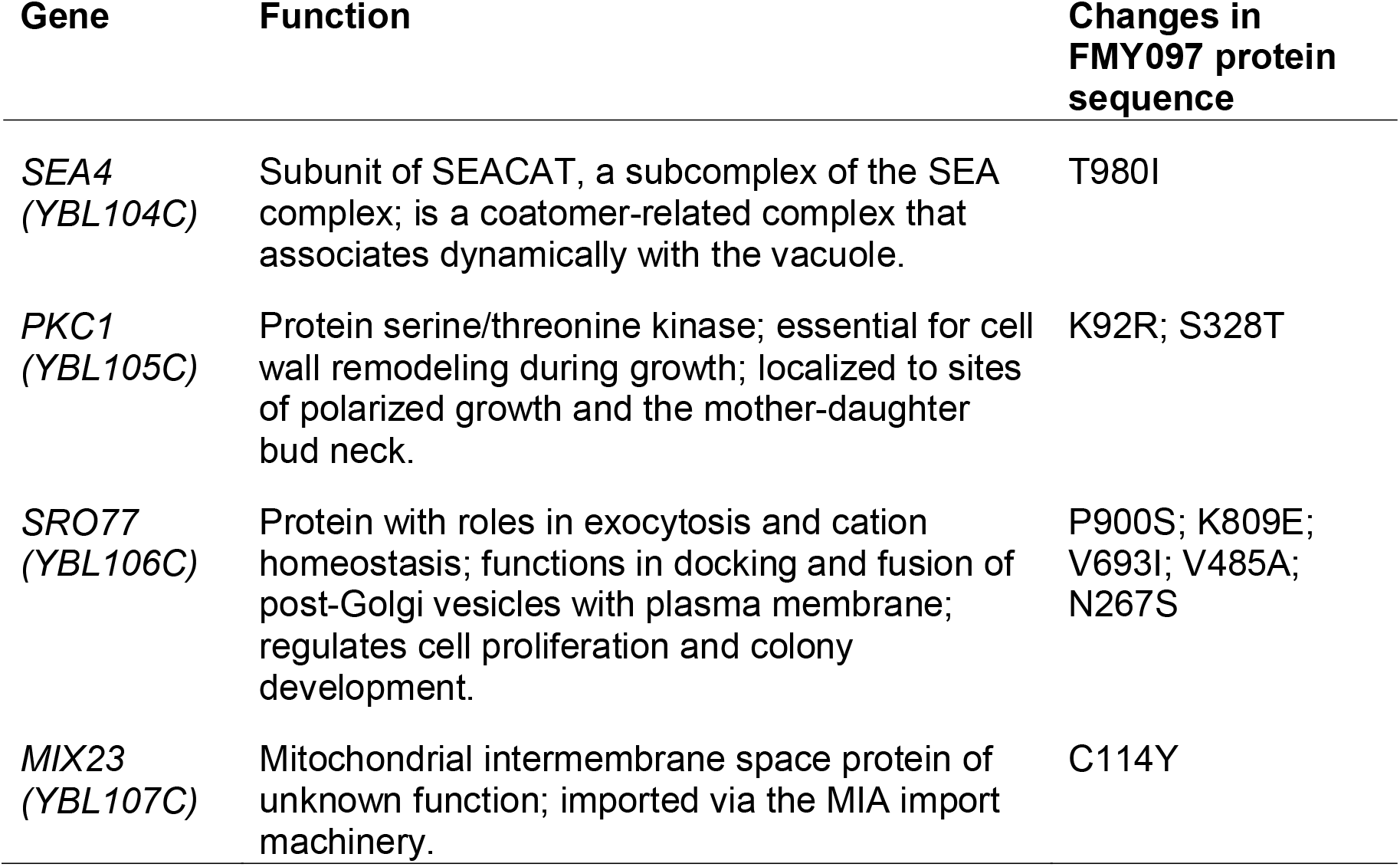
HMF resistance-related candidate genes in chromosome II.

### 3.4. VALIDATION OF CAUSATIVE GENES IN THE HMF RESISTANCE PHENOTYPE

Following the QTL mapping, the contribution of alleles *MIX23*, *PKC1*, *SEA4,* and *SRO77* to the HMF resistance phenotype was performed using the traditional reciprocal hemizygosity analysis (RHA) approach. In short, a knocked-out version of each allele of either parental of the hybrid FMY097/BY4742 is crossed with the wild-type version of the other parental. It is important to note that *PKC1* is an essential gene and null mutants can only be recovered in the presence of an osmotic stabilizer (**see Materials and Methods for details**). Despite the effort of an extensive screening for a *pkc1*Δ version of strain FMY097, such an event was not observed. On the other hand, a knocked-out version of this allele in BY4742 was accomplished. Nevertheless, RHA of the alleles *MIX23*, *SEA4,* and *SRO77* did not show loss of robustness in any mutated versions of the hybrid (**see Supplementary Fig S4**), indicating that none of these genes is a major contributor to the evaluated phenotype. From this assay, we could also observe that resistance to HMF is a dominant trait in FMY097.

Next, because we ought to understand the effect of *PKC1*^FMY097^ in HMF resistance, we cloned this allele in strain BY4742 *pkc1*Δ. To complete the analysis, and to investigate if the other genes could have a minor effect on the phenotype, we performed the same genetic transformation for the other 3 alleles - *MIX23*^FMY097^, *SEA4*^FMY097^, and *SRO77*^FMY097^. The overall fitness of mutated BY4742 strains in control conditions did not change, except for strain’s BY4742 *PKC1*^FMY097^ lower maximum specific growth rate (**see Supplementary Fig S5**). All the modified versions of the susceptible strain BY4742 were phenotyped in presence of HMF, both in solid and liquid media - the former to infer subtle response changes in mutants that cannot be measured in traditional spot assays. An intermediate concentration of 10 mM HMF was also tested to infer genetic contributions to mild stress. The results are depicted in Fig 3 and all the kinetic parameters extracted from the growth curves are displayed in Table 2.

**Fig 3:**
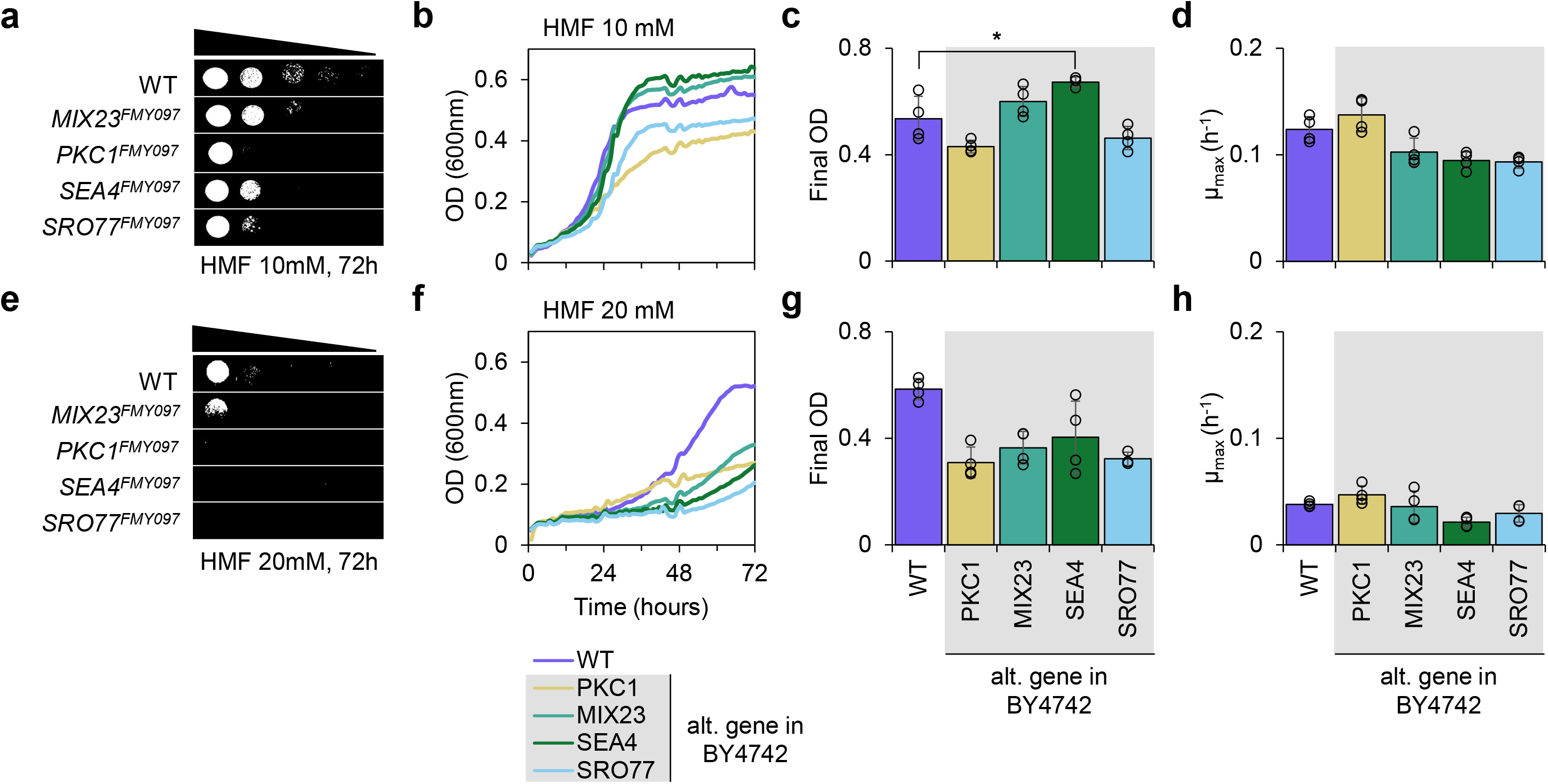
Phenotyping in HMF of strain BY4742 with FMY097 alleles found in the QTL window of HMF resistance genetic mapping. WT stands for the wild-type version of BY4742. **a to d:** (top half of the image) represent phenotyping in 10 mM HMF; **e to h:** (bottom half of the image) represents phenotyping in 20 mM HMF. **a and e:** spot test in HMF after 72h incubation of wild-type (WT) and modified versions of BY4742 with *MIX23*^FMY097^, *PKC1*^FMY097^, *SEA4*^FMY097^ and *SRO77*^FMY097^. **b and f:** representative growth curves of WT and modified versions of BY4742 with alternative (alt.) genes from FMY097 (lines represent the average of 4 replicates). **c and g:** bar chart of the final OD and **d and h:** of the maximum specific growth rate of WT and modified versions of BY4742, respectively. Black circles in the bar charts represent the replicate data. (*) stands for p-value < 0.05

As previously mentioned, HMF greatly impairs BY4742 fitness, and this statement is confirmed in Fig 3. When 10 mM HMF is present, the alternative FMY097 alleles do not endow robustness in cell growth in solid media (Fig 3a), and one could say that the mutated genes harmed BY4742 intrinsic robustness in the spot test. On the other hand, growth in liquid media (Fig 3b) had a different result, indicating that different mechanisms of HMF tolerance act in distinct cultivation status. In microplate cultivation, a higher final OD was obtained by the strains harboring mutated *MIX23* (0.600 ± 0.057) and *SEA4* (0.672 ± 0.020), compared to the wild-type BY4742 (0.535 ± 0.084) (Fig 3c). In this scenario, *SEA4*^FMY097^ significantly improved final OD when growing in 10 mM HMF. Meanwhile, *SEA4*^FMY097^ and *SRO77*^FMY097^ reduced the strain’s maximum specific growth rate (Fig 3d). In harsher conditions, however, alternative alleles have depleted BY4742 cell growth in either solid (Fig 3e) and liquid (Fig 3f) media. 20 mM HMF had negative effects in final OD (Fig 3g) of BY4742 modified strains, but no outcome in the strains’ µ_MAX_ (Fig 3h). Deleterious effect in mutated BY4742 reveals a strain-specific correlation to the phenotype, possibly caused by a background effect in FMY097.

### 3.5. GENETIC NETWORK AND THE REDOX ENVIRONMENT IN FMY097

Given that the most prominent region in the QTL mapping analysis revealed a causative allele with minor contribution to HMF resistance (*i.e., SEA4*^FMY097^ was able to improve BY4742 growth in presence of this aldehyde but did not fully recover FMY097’s fitness), epistatic or additive effects are expected. The transfection of FMY097’s complete *MIX23-SEA4* genomic region to BY4742 did not result in transformants - the presence of an essential gene might have hampered the process. Because there is evidence that *PKC1* and *SRO77* interact genetically (Liou et al. 2014) and are upregulated in presence of HMF (Zhou et al. 2014; Qiu and Jiang 2017), strain BY4742 *PKC1*^FMY097^ *SRO77*^FMY097^ was constructed. Nevertheless, no improvement in HMF resistance was observed (results not shown). Testing of other FMY097 allele combinations in BY4742 would result in a labor-intensive genetic engineering effort and was not performed.

Next, we proceeded to evaluate the *in silico* interaction network arising from *MIX23*, *PKC1*, *SEA4,* and *SRO77* (Fig 4). A total of 20 other genes and 516 possible links (mainly physical and genetic interactions) are present in this network, representing a complex genetic architecture (Fig 4a). Gene ontology of this cluster reveals that the main processes involved are related to the regulation of TORC1/TOR signaling, general intracellular signal transduction and organelle assembly (Fig 4b) (**see Supplementary Table S4**).

**Fig 4:**
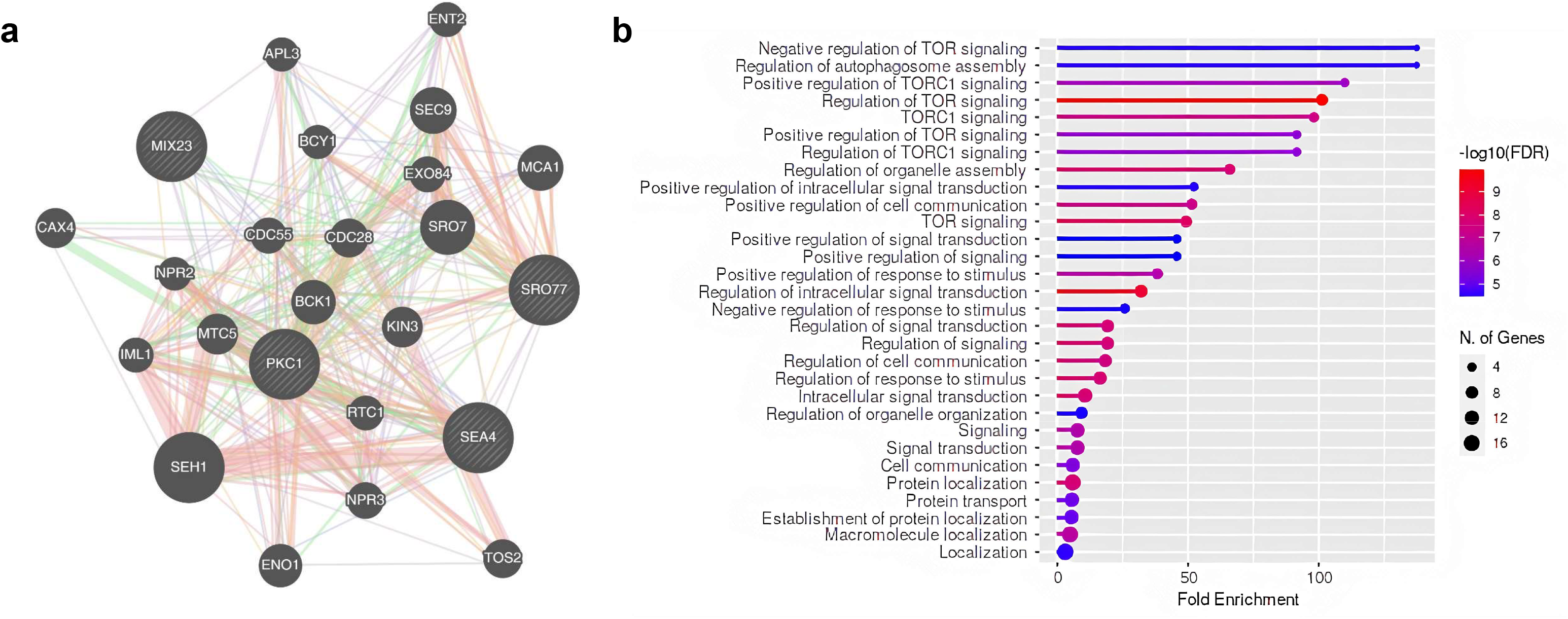
*In silico* analysis of the interaction between alleles present in QTL mapping of HMF resistance in strain FMY097. **a:** Genetic network of *MIX23*, *SRO77*, *SEA4*, and *PKC1* (performed in GeneMania (Warde-Farley et al. 2010)). Physical interaction (red lines): two gene products are linked if they were found to interact in a protein-protein interaction study; genetic interaction (green lines): two genes are functionally associated if the effects of perturbing one gene were found to be modified by perturbations to a second gene. **b:** Gene ontology enrichment analysis of the biological processes involved in the 24 genes network, as performed in ShinyGO v0.741 (Ge et al. 2020)

Because most genes implicated in endogenous detoxification reactions of aldehydes such as HMF and furfural - both present in lignocellulosic hydrolysate - are NADPH-dependent (Wang et al. 2018) (Fig 5a), we moved on to test whether the intrinsic cofactor availability in the parental strains of hybrid FMY097/BY4742 could contribute to understanding the differences in the HMF tolerance phenotype. The quantification of this cofactor was performed using an indirect luminescence assay in optimal growth conditions. Results show that FMY097 has outstandingly more NADPH than BY4742 (p-value = 0.002) (Fig 5b) and, therefore, a favorable background for reduction reactions where this cofactor takes place. Amongst other possible causes, NADPH availability can indeed elucidate the HMF resistance in FMY097.

**Fig 5:**
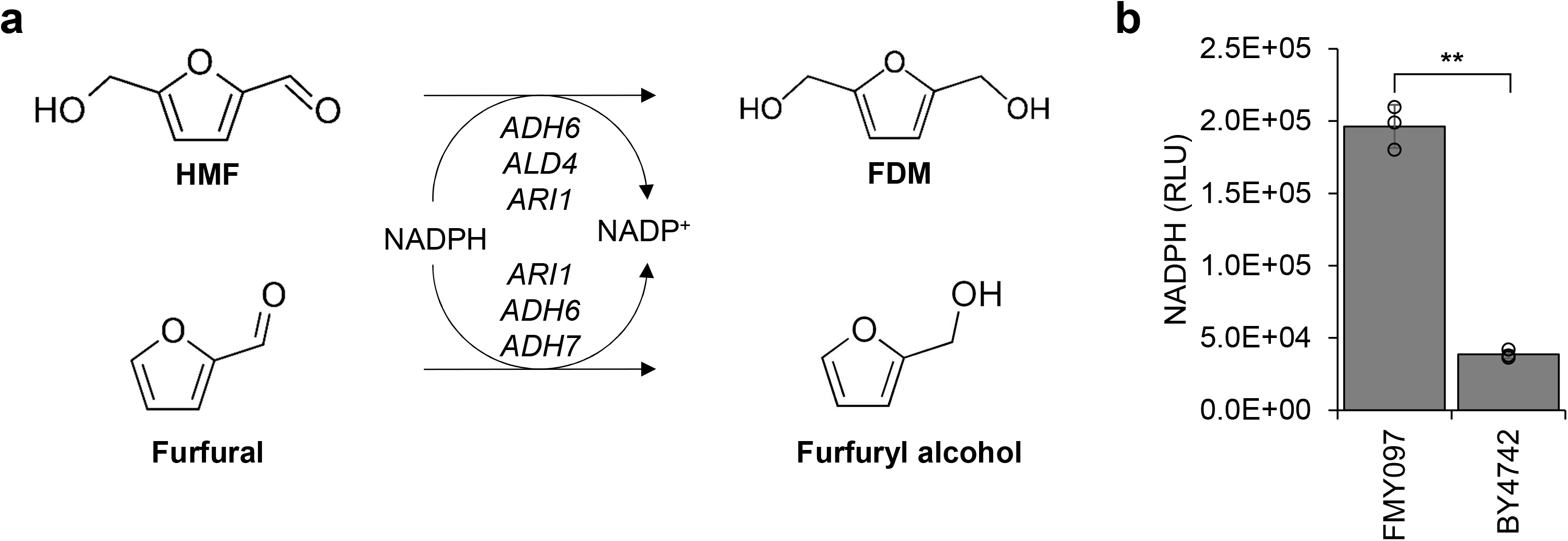
NADPH quantification in strains FMY097 and BY4742. **a:** Main endogenous reaction in *S. cerevesiae* for the detoxification of HMF and furfural. The aldehyde and alcohol dehydrogenase enzymes-encoding genes shown are NADPH-dependent. **b:** NADPH as a measure of Relative Luminesce Units (RLU) of strains FMY097 and BY4742. Black circles in the bar chart represent the replicate data. (**) stands for p-value < 0.01

### 3.6. CANDIDATE GENES CONTRIBUTION TO THERMOTOLERANCE

As a follow-up to the examination of the candidate alleles’ contribution to HMF tolerance, we tested the mutated BY4742 strains’ performance in heat stress. This decision was based on the thermotolerant background of parental strain FMY097 and the fact that the essential gene *PKC1* - part of the CWI pathway, described as participating in heat response (Auesukaree et al. 2009) - was assigned as a potential contributor in the HMF QTL mapping with uncommon non-synonymous mutations. The phenotyping in high temperature was performed in the same fashion as previously stated for the HMF resistance, and the results are shown in Fig 6 and Table 2.

**Fig 6.**
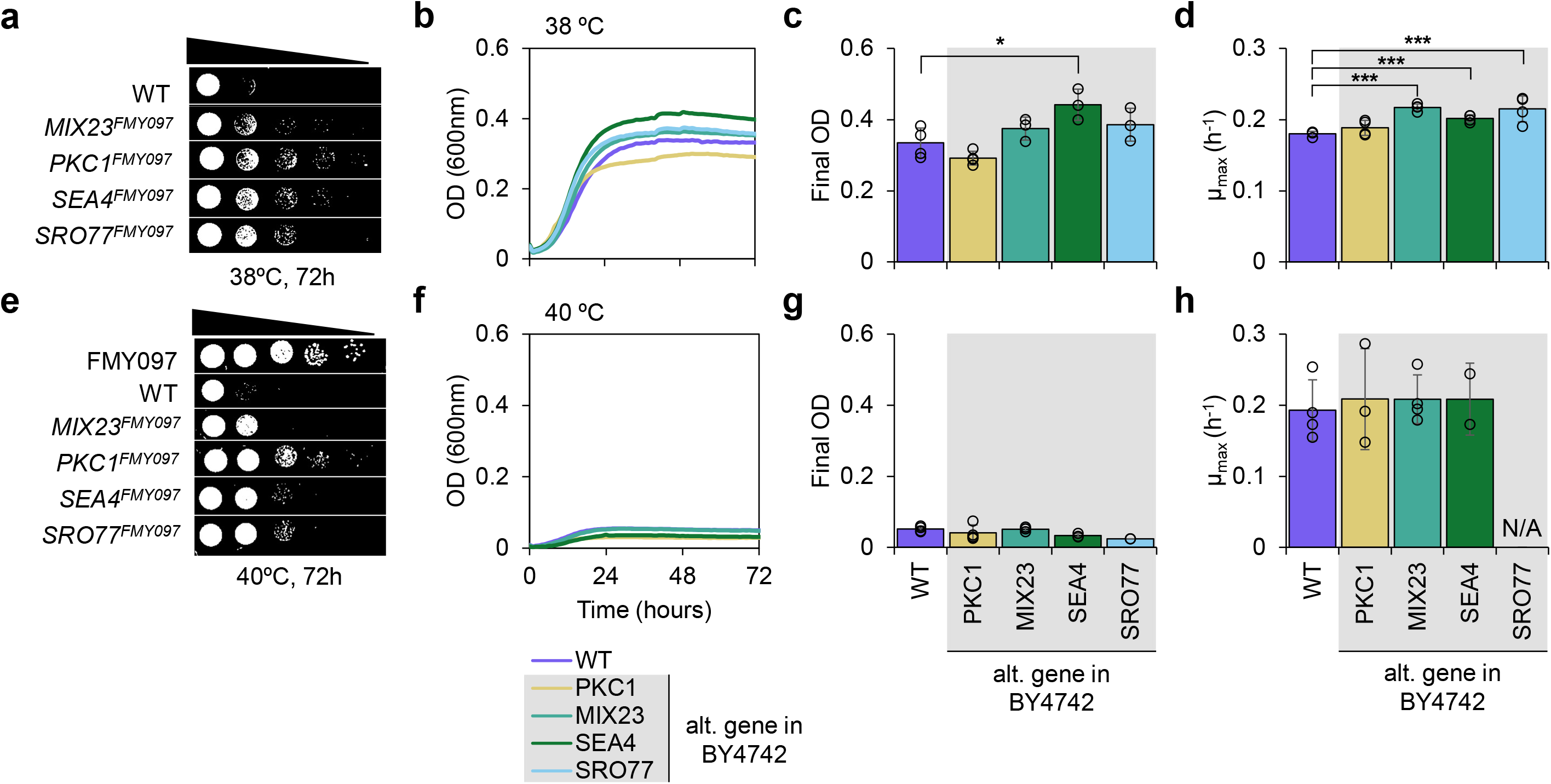
Phenotyping in high temperature of strain BY4742 with FMY097 alleles found in the QTL window of HMF resistance genetic mapping. WT refers to the wild-type version of BY4742. **a to d** (top half of the image) represent phenotyping in 38 °C; **e to h** (bottom half of the image) represent phenotyping in 40 °C. **a and e:** spot test after 72h incubation of wild-type (WT) and modified versions of BY4742 with *MIX23*^FMY097^, *PKC1*^FMY097^, *SEA4*^FMY097^ and *SRO77*^FMY097^; FMY097 is also shown in image **e**. **b and f:** representative growth curves of WT and modified versions of BY4742 with alternative (alt.) genes from FMY097 (lines represent the average of 4 replicates). **c and g:** bar chart of the final OD and **d and h:** of the maximum specific growth rate of WT and modified versions of BY4742, respectively. Black circles in the bar charts represent the replicate data. (*) stands for p-value < 0.05 and (***) for p-value < 0.001

Similar results obtained in the HMF tolerance assay were also observed for heat stress response. At 38 °C, strain BY4742 has a 52% loss in cell density after 72h cultivation, while for FMY097 this number is 33%. When phenotyped in solid media (Fig 6a), it is visible that all the mutated versions of BY4742 bearing FMY097 alleles improved the overall performance compared to the unmodified strain. Here, the genes *MIX23, PKC1,* and *SEA4* with FMY097 mutations particularly presented the best results. Growth in liquid media (Fig 6b) had different results, as it was observed for HMF resistance. Final OD of BY4742 *SEA4*^FMY097^ (0.442 ± 0.044) was significantly superior to the wild-type (WT) version (0.334 ± 0.042) (Fig 6c). The maximum specific growth rate of strains BY4742 *MIX23*^FMY097^, BY4742 *SEA4*^FMY097^, and BY4742 *SRO77*^FMY097^ improved at 38 °C (Fig 6d). When at 40 °C, growth in solid media (Fig 6e) had an important outcome: strain BY4742 *PKC1*^FMY097^ was able to produce cells at the lowest dilutions after 72h cultivation, outperforming the other BY4742 versions. It is possible to notice that the mutated *PKC1* in BY4742 almost fully recovered the industrial parental FMY097 response to this temperature challenge. Growth in liquid media (Fig 6f), on the other hand, did not present the same result and any strain was able to grow.

### 3.7. PREVALENCE OF *SEA4* AND *PKC1* FMY097 MUTATIONS

Because the FMY097 mutated versions of *SEA4* had positive effects in mild aldehyde and temperature challenges in liquid media and *PKC1* at high-temperature cultivation in solid media, an in-depth analysis of such protein sequences change is deemed necessary. For that, the sequence of proteins Sea4p and Pkc1p were aligned to 1,015 other relevant *S. cerevisiae* strains (Fig 7). The main bioethanol *S. cerevisiae* strains (PE-2, CAT-1, BG-1 and Ethanol Red) protein sequences were obtained from (Nagamatsu et al. 2021) and the other 1,011 were available in (Peter et al. 2018)

**Fig 7.**
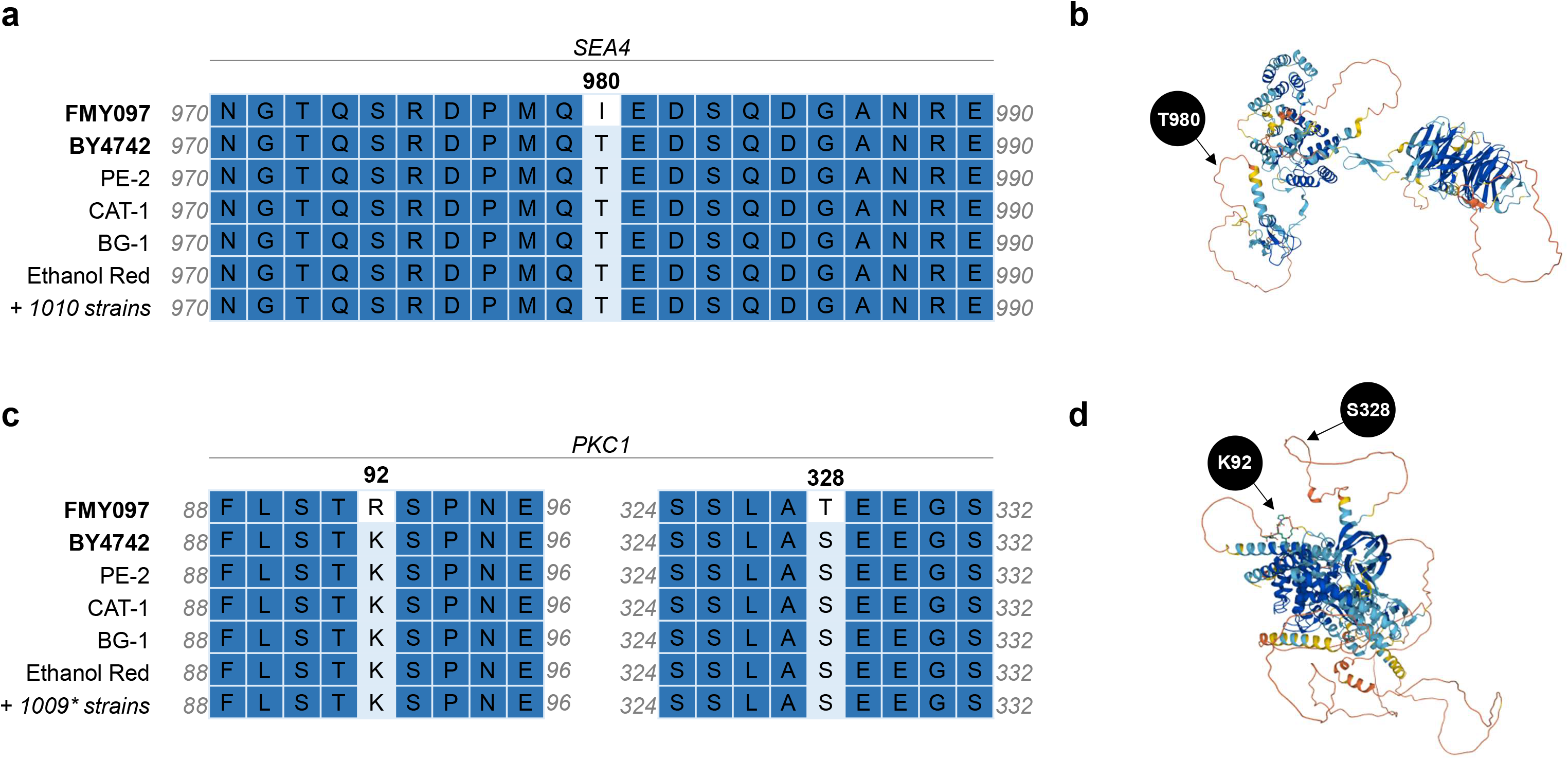
Analysis of mutations present in *SEA4*^FMY097^ and *PKC1*^FMY097^. **a:** Protein sequence alignment surrounding position 980 in Sea4p between strains FMY097 and other relevant *S. cerevisiae* strains. **b:** Sea4p three-dimensional protein structure with an indication of amino acid T980. **c:** Protein sequence alignment surrounding position 92 and 328 in Pkc1p between strains FMY097 and other relevant *S. cerevisiae* strains. **d:** Pkc1p three-dimensional protein structure with an indication of amino acid K92 and S328. *For this alignment, the number of other *S. cerevisiae* is 1009 for mutation S328T and 1010 for mutation K92R

The T980I mutation in Sea4p does not appear in other typical bioethanol industrial strains and 1,010 other sequences evaluated, indicating a very uncommon specific mutation for FMY097 (Fig 7a) - a frequency of 0.2% in *S. cerevisiae*. The only strain with the same mutation amongst the 1,011 is denominated CBS7959 - an isolate from the Brazilian sugarcane ethanol industry, the same environment where SA-1 (FMY097 parental) is derived from. The non-synonymous mutation is located in a connector region in the protein - namely loop (Papaleo et al. 2016) -, outside of known domains (Fig 7b). As for the two mutations present in Pkc1p, the results are similar to the mutations in *SEA4* - not found in the main industrial and other relevant *S. cerevisiae* (Fig 7c). Mutation K92R is also present in CBS7959, representing a frequency of 0.2%. Meanwhile, besides CBS7959, mutation S328T can also be found in strain CEY647 - isolated from *Molussus molussus* (bat), resulting in a 0.3% frequency. Protein modifications in *PKC1* are also very rare, located in loop regions that do not contain a domain (Fig 7d). Mutation S328 is localized in a disordered region composed of polar residues. Altogether, the NS mutations in FMY097 that can induce better performance in lignocellulosic hydrolysates are very rare.

## 4. DISCUSSION

Understanding the genetic basis of complex traits related to industrial *Saccharomyces cerevisiae* robustness is a great challenge for improving ethanol productivity. Adverse conditions present during hydrolysate fermentation in the second-generation industry are diverse and include thermal stress and metabolism inhibitors, such as aldehydes. One of these compounds, HMF, has a rather destructive power in yeasts, compromising cell viability. While transcriptomic analysis has been performed in yeast subjected to this fermentation inhibitor, QTL mapping of HMF tolerance is an important step towards unraveling mechanisms of superior aldehyde resistance in *S. cerevisiae* for reverse engineering endeavors. In this work we used a hybrid Bulk (BSA) and Individual (ISA) Segregant Analysis for this purpose, where haploids were individually phenotyped and bulk sequenced.

Firstly, a heterozygous hybrid for HMF resistance was built. The previously described highly aldehyde tolerant haploid FMY097 shortened the process for QTL analysis. The crossing with laboratory BY4742 was decided given its well-known genotype and documented susceptibility towards HMF (Zhou et al. 2014). Obtaining large pools of FMY097/BY4742 segregants was performed by specific mating type fluorescence detection in flow cytometry cell sorting. The low efficiency in collecting MATα cells – only 8% of total collected individuals had the correct mating type – can be explained by the equipment insensibility in detecting orange fluorescence. Nevertheless, the ∼500 FMY097/BY4742 segregant library was large enough for further screening of two pools with 60 haploids with opposite HMF resistance phenotype. Sequencing a large population of organisms with distinct extreme phenotypes is crucial for precising the mutations in QTL, given that it improves statistical analysis of genetic markers (Wilkening et al. 2014). It is important to note that the population colony size distribution in 20 mM HMF was normal, typical of quantitative traits. BSA of the 120 segregants of hybrid FMY097/BY4742 identified a Δ(SNP-index) peak region in chromosome II containing 4 genes with at least one non-synonymous uncommon SNP in FMY097: *SEA4*, *PKC1*, *SRO77* and *MIX23*.

Out of the 4 genes implicated in the genomic region related to superior HMF resistance, *PKC1* and *SRO77* have been previously associated with response to this aldehyde. *PKC1* is part of the three-tiered cascade of the mitogen-activated protein kinase (MAPK) pathway, that regulates cell proliferation, differentiation, survival, and death (Herskowitz 1995; Chen and Thorner 2007). Zhou et. al (2014) (Zhou et al. 2014) revealed that *PKC1* is upregulated in the CWI and high osmolarity glycerol (HOG) of the MAPK pathway in response to challenges of 30 mM HMF. The group also reported that *PKC1* showed consistent enhanced signature expression in the phosphatidylinositol signaling pathways, which mediate numerous physiological processes, in response to the aldehyde. Furthermore, Vilella et. al (2005) (Vilella et al. 2005) stated that overexpression of *PKC1* increases cell viability under oxidative stress - which effects on yeasts are similar to HMF - because it enhances the machinery required to repair the altered cell wall and restore actin cytoskeleton polarity by promoting actin cable formation. Meanwhile, S*RO77* (*SRO7* homologue), involved in docking and fusion of post-Golgi vesicles with plasma membrane, was found upregulated after 12h of very high gravity fermentation with 17.5 mM HMF in an engineered yeast strain with improved ethanol tolerance and productivity (Qiu and Jiang 2017).

QTL mapping usually reveals a major causative gene to a given phenotype in yeast (Feng et al. 2018; Coradini et al. 2021), but it was not the case for this study. Here, typical RHA was not able to uncover one major genetic contribution to HMF resistance and, therefore, allele interchange deemed necessary. This is an important note on QTL studies, given that minor endowment in robustness can represent important process optimization, emphatically in industrial applications. Therefore, BY4742 mutants bearing *SEA4*, *PKC1*, *SRO77* and *MIX23* alleles from FMY097 were tested in presence of HMF. While no improvement in phenotype was observed at 20 mM HMF, BY4742 *SEA4*^FMY097^ was able to produce more cell density in liquid media containing 10 mM HMF compared to the non-edited laboratory strain. There is no available data on the profile expression of *SEA4* in the presence of this aldehyde, and this result configures as the first report on the influence of this allele to HMF resistance.

*SEA4* integrates the SEA (Seh1-associated) protein complex that dynamically associates with the vacuole *in vivo* and has a role in intracellular trafficking, amino acid biogenesis, and response to nitrogen starvation (Dokudovskaya et al. 2011). The SEh1-Associated protein 4 is part of the SEACAT (SEA subcomplex activating TORC1) that is an upstream regulator of TORC1 (Target of Rapamycin Complex 1) pathway, involved in various cellular activities, such as cell growth, ribosome biogenesis, translation initiation, metabolism, stress response, aging, and autophagy (Inoue and Nomura 2018). TORC1 integrates signals from a wide variety of intracellular and extracellular cues to maintain optimal growth and metabolism (Dokudovskaya and Rout 2015; Loissell-Baltazar and Dokudovskaya 2021) and has been reported to sense and respond to oxidative stress in a complex manner (Su and Dai 2017). Furthermore, *PKC1* is a direct substrate of TORC2 (Nomura and Inoue 2015)( - another complex of the TOR kinase - that is also activated in response to oxidative stress in *S. cerevisiae* (Niles and Powers 2014). Altogether, these findings corroborate with the GO enrichment analysis performed with the genetic network arising from the QTL, in which *PKC1* and *SEA4* are involved in notorious intracellular signaling pathways.

Besides granting results for the genotype-phenotype relationship between point genetic mutations and performance in media containing HMF, allele interchange also provided insight into gene-gene interaction for this trait in FMY097. Although data from empirical studies of genetic variance components imply that additive variance typically accounts for over half to 100% of the total genetic variance (Hill et al. 2008), recent study have shown that non-additive effects of gene-background interactions can’t be neglected in *S. cerevisiae* (Sardi and Gasch 2018). Sardi and Gash (Sardi et al. 2018), investigating mechanisms of tolerance to lignocellulosic toxins found in plant hydrolysate across 165 *S. cerevisiae* lineages, found that most of the implicated genes played strain-specific roles in inhibitor tolerance. Thus, whether the gene is required for toxin resistance in a given background can play a bigger role than the gene’s allele. In this study, minor allelic contribution added to the deleterious effects in mutated BY4742 strain at 20 mM HMF indicates a substantial interaction of the implicated genes with the genetic background.

While the complex genetic network resulting from the QTL mapping of FMY097 HMF resistance is mainly related to intracellular signaling - in specific for TORC1 and CWI pathways -, another global feature that can interfere in aldehyde detoxification, specially HMF and furfural, is NADPH availability. In fact, such feature has traditionally played a key part in enhancing aldehyde tolerance in *S. cerevisiae* by either overexpressing (i) NADPH-dependent alcohol or acetaldehyde reductases to promote the conversion to less-toxic products (Moon and Liu 2012; Hasunuma et al. 2014) (REF); or (ii) genes related to the interconversion of NADPH and NADP^+^ (Liu et al. 2020). While cofactor engineering can create better redox environments for more resistant yeast strains, here we show that HMF-tolerant FMY097 presents an intrinsic favorable NADPH environment. This result suggests that indigenous industrial strains subjected to high productivity efforts can develop improved intracellular redox levels that lead to inhibitor robustness and represent a useful chassis to produce reduced sugars.

Alternative alleles found in FMY097 were also tested for contribution to heat stress tolerance. As previously mentioned, this decision was based on the different membrane functions of genes found in the HMF resistance QTL mapping and their participation in major intracellular signaling pathways - in special *PKC1* and CWI, already associated with thermotolerance (Auesukaree et al. 2009) - and the outstanding ability of FMY097 to grow at 40 °C. Again, *SEA4*^FMY097^ was able to enhance BY4742 fitness at elevated temperature in liquid medium, indicating that TORC1 is an important regulator of cell response to different stress. The outcome of mutated BY4742 phenotyping in solid medium was different: the laboratory strain expressing *PKC1*^FMY097^ outperformed the other in temperatures as high as 40 °C, fully recovering the industrial parental growth. Not only does this result confirm the importance of the MAPK regulator *PKC1* in thermotolerance, but also provides an unprecedented contribution to one of the most studied yeast stress factors. It is important to note that the cultivation status greatly influenced the strain’s phenotype, as differences in features such as oxygen supply can affect biochemical reactions and activate different gene clusters (Pan et al. 2019). This is an indication that a comprehensive and diverse screening is paramount for genotype-phenotype association assays.

Finally, the frequency of the NS mutations in *SEA4*^FMY097^ and *PKC1*^FMY097^ were calculated and were found to be very rare. Amino acid changes T980I in *SEA4* and K92R and S328T in *PKC1* are all located in connector regions in the protein. These regions, known as loops, connect secondary structure elements and concentrate the higher number of single nucleotide variations in the surface of proteins (Bhattacharya et al. 2017). Surface-exposed loops are responsible for important functional roles contributing to the formation of binding and catalytic sites (Papaleo et al. 2016), playing a wide range of protein functions, such as interaction to other proteins in signaling cascades (Zomot and Kanner 2003; Bernstein et al. 2004). Together, this evidence indicates that the almost-exclusive FMY097’s protein mutations might contribute to better protein-protein binding (as in Sea4p to the SEA complex) or more effective signal transduction (as in Pkc1p in the CWI or Sea4p in the TORC1 pathways).

Unlike previous genome-wide association studies that suggested that HMF resistance in yeast could be based on the overexpression of reductases or dehydrogenases, our results suggest that a tolerant strain might have mutations in genes that have important roles in different membrane functions and regulators of cell growth and stress response pathways. As described by Liu *et. al* (Liu et al. 2018), tolerant industrial yeast strains possess a more robust CWI pathway signaling against HMF and our results corroborate this finding. Also, the description of *SEA4* as a global participant in important industrial stress response adds to the understanding of the complex TOR mechanisms in yeast.

## Supporting information

Supplementary Images

## AUTHOR’S CONTRIBUTIONS

FSBM conducted all experiments and wrote the manuscript. ALVC provided important discussion and support throughout this study. MFC analyzed genomic data. CM contributed to the genetic engineering of the strains. MF developed the plasmid used for segregant collection in flow cytometry. GAGP and GST conceived and supervised the project. All authors read and approved the final manuscript.

## ETHICS APPROVAL AND CONSENT TO PARTICIPATE

Not applicable.

## CONSENT FOR PUBLICATION

Not applicable.

## AVAILABILITY OF DATA AND MATERIAL

The complete dataset of DNA-seq reads have been deposited in SRA under accession number PRJNA781913.

## ADDITIONAL FILES

**Additional File 1**

**Format:** Document (DOCX)

**Title: Supplementary Images**

**Description:** Supporting images described in this study.

**Additional File 2 Format:** Table (XLS)

**Title: Supplementary Tables**

**Description:** Supporting tables described in this study.

