## Supplementary Images for "Genetic mapping of a bioethanol yeast strain reveals new targets for aldehyde- and thermotolerance"

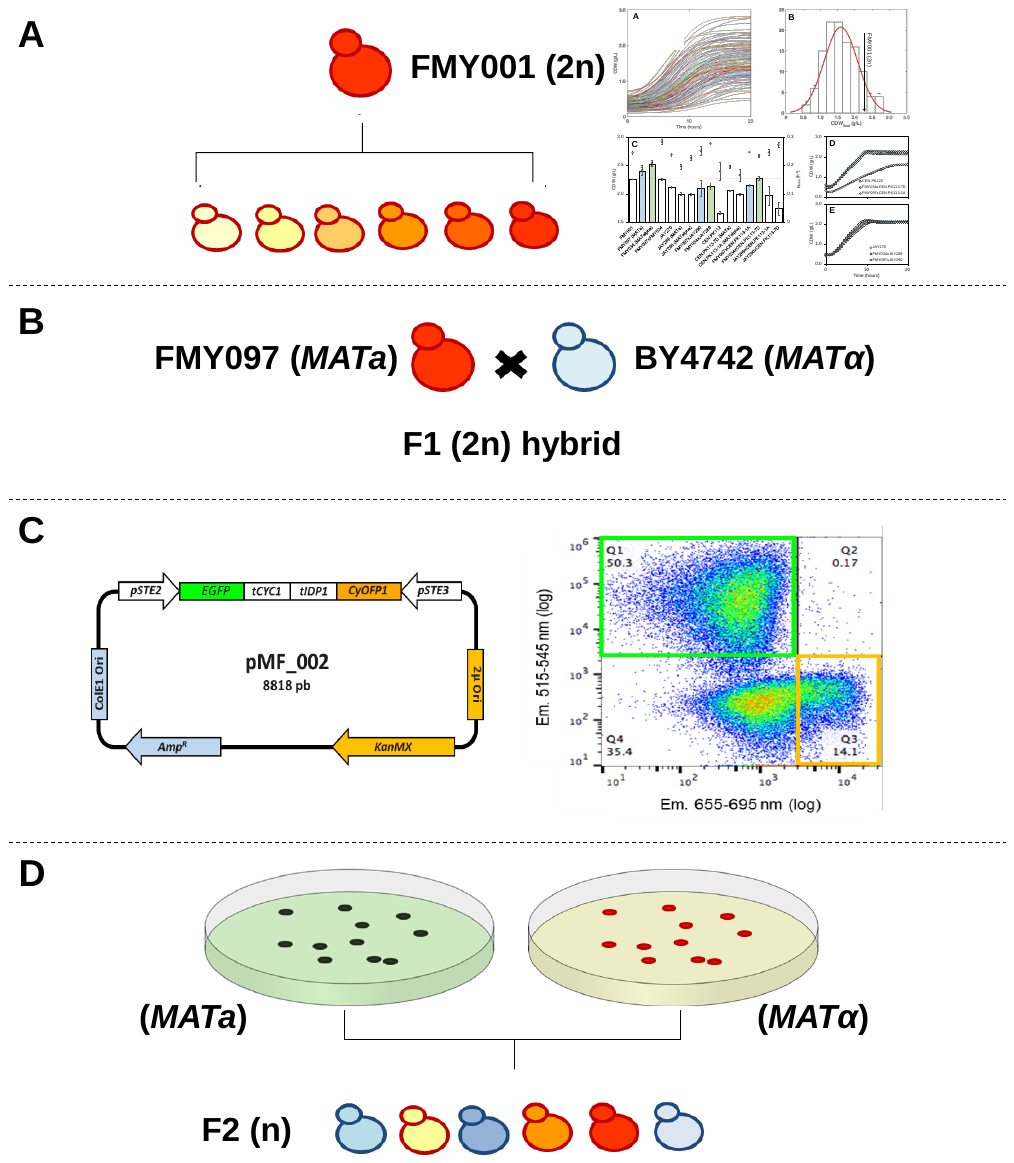

**Fig S1** Graphic representation of the methodology used for the obtaining of haploids progeny of the crossing FMY097/BY4742 for QTL mapping of HMF resistance. (**A**) Screening of the FMY001 (2n) segregants using individual segregant analysis (ISA) phenotyping for the selection of a highly HMF resistant haploid, using methods described by de Mello *et al* (2019); (**B**) Construction of a highly heterozygous hybrid from the crossing of FMY097 and BY4742. (**C**) Plasmid pMF002 used in the transformation of the hybrid FMY097/BY4742 (left) and graphic of the gates used in flow cytometer to collect segregants (right). (**D**) Obtaining haploids with high genetic variability for the HMF resistance phenotype using a cell sorter.

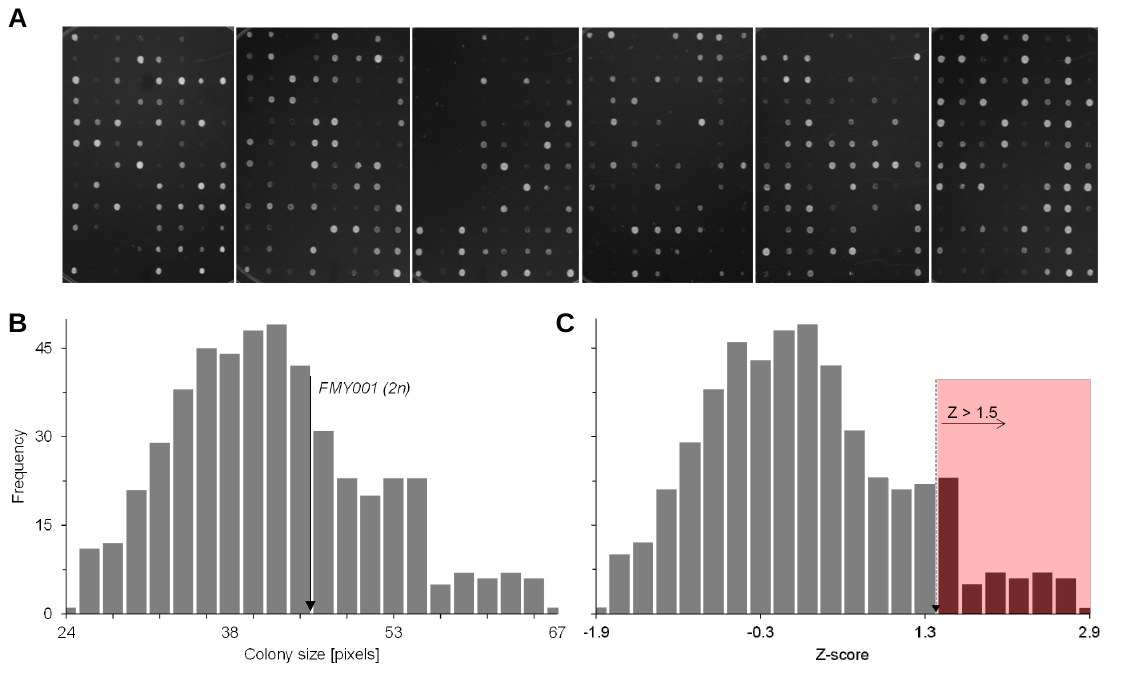

**Fig S2** Phenotypic analysis of the 494 FMY097/BY4742 segregants. (**A**) Photos of the 6 plates after 48h of growth SC + 20 mM HMF. These images were used in the software ImageJ for determining colony size, in pixels, of each haploid. Bigger colonies (more pixels) represent strains resistant to HMF; smaller colonies (less pixels) represent strains susceptible to the inhibitor. (**B**) Segregants colony size distribution histogram, with a marker (arrow) for the size of the original parental strain FMY001 (2n). (**C**) Z-score distribution histogram for each segregant, highlighting the ones with values over 1.5.

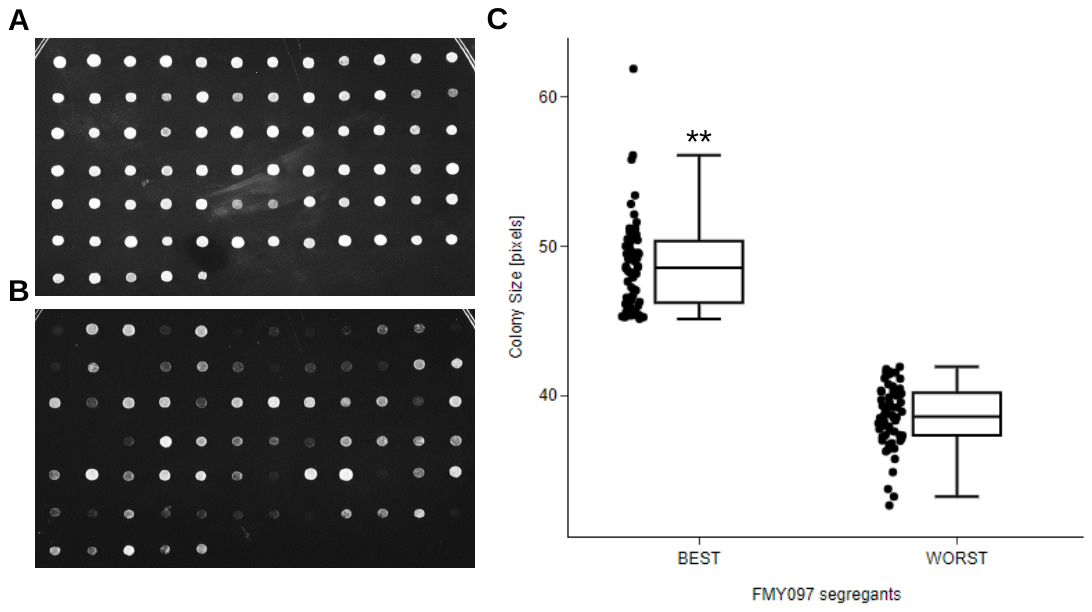

**Fig S3** Selection of the segregant pools with extreme phenotypes for sequencing and QTL mapping. (**A**) Photo of the 60 superior segregants after 48h of growth in 20 mM HMF. (**B**) Photo of the 60 inferior segregants after 48h of growth in 20 mM HMF. (**C**) Boxplot of the colony size distribution of the *BEST* and *WORST* pools derived from the parental FMY097/BY4742 (**) p-value < 0.0001.

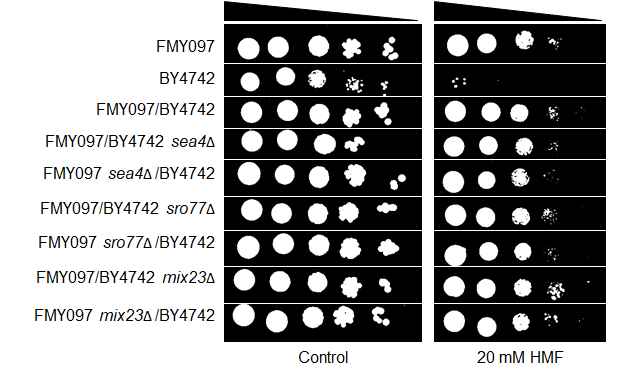

**Fig S4** Reciprocal hemizygosity analysis of candidate genes in the QTL mapping of HMF resistance. Image shows a spot test of 10x dilutions (starting at OD_600_ = 1) for the hemizygotes of alleles *SEA4, MIX23,* and *SRO77* in FMY097 and BY4742 taken after 72h cultivation. *On the left*: control condition in SC medium; *on the right*: SC + 20 mM HMF medium.

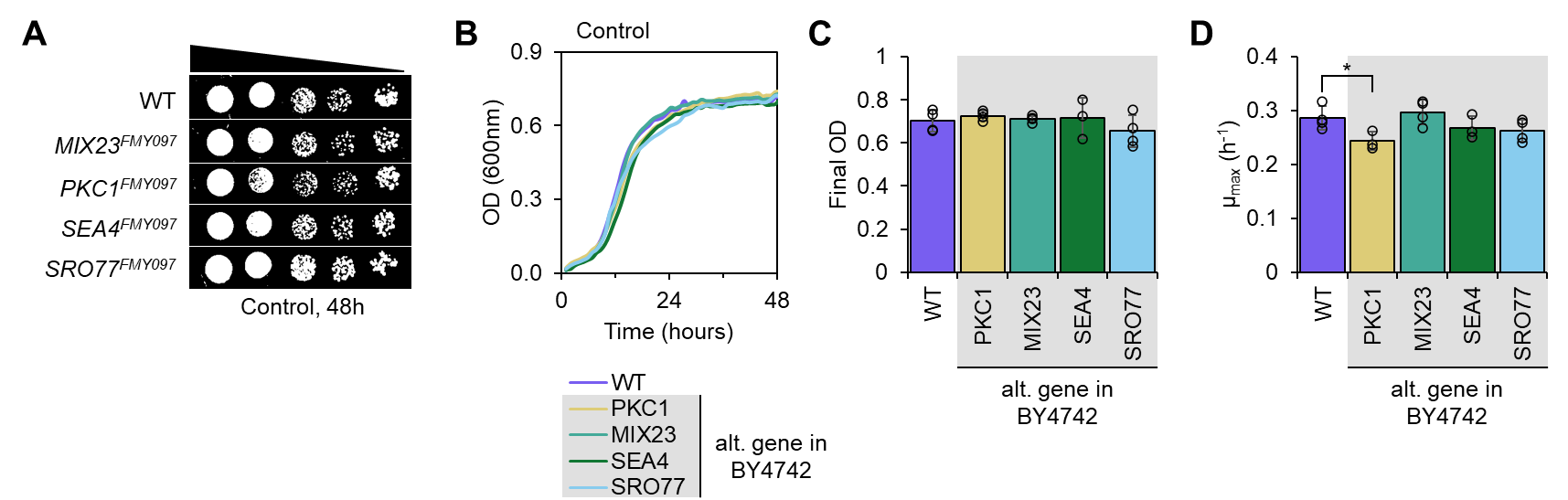

**Fig S5** Phenotyping in SC medium of strain BY4742 with FMY097 alleles found in the QTL window of HMF resistance genetic mapping. WT stands for the wild-type version of BY4742. **A**: spot test in SC after 48h incubation of wild-type (WT) and modified versions of BY4742 with *MIX23*^FMY097^, *PKC1*^FMY097^, *SEA4*^FMY097^ and *SRO77*^FMY097^. **B**: representative growth curves of WT and modified versions of BY4742 with alternative (alt.) genes from FMY097 (lines represent the average of 4 replicates). **C**: bar chart of the final OD and **D:** of the maximum specific growth rate of WT and modified versions of BY4742, respectively. Black circles in the bar charts represent the replicate data. (*) stands for p-value < 0.05.
